# Dot1L-dependent H3K79 methylation facilitates histone variant H2A.Z exchange at DNA double strand breaks and is required for high fidelity, homology-directed DNA repair

**DOI:** 10.1101/544981

**Authors:** Nehemiah S. Alvarez, Pavla Brachova, Timothy A. Fields, Patrick E. Fields

## Abstract

In eukaryotic cells, the homology-directed repair (HDR) and non-homologous end joining (NHEJ) pathways are required for the repair of DNA double strand breaks (DSB). The high-fidelity HDR pathway is particularly important for maintenance of genomic stability. In mammals, histone post-translational modifications and histone variant exchange into nucleosomes at sites of DSB generate an open chromatin state necessary for repair to take place. However, the specific contributions of histone modifications to histone variant exchange at DSB sites and the influence of these changes on the DNA repair process and genome stability are incompletely understood. Here we show that Dot1L-catalyzed methylation of H3 histone on lysine 79 (H3K79) is required for efficient HDR of DSB. In cells with DNA DSB either lacking Dot1L or expressing a methylation-dead Dot1L, there is altered kinetics of DNA repair factor recruitment, markedly decreased H2A.Z incorporation at DSB sites, and a specific and profound reduction in HDR, which results in significant genomic instability. These findings demonstrate a new role for Dot1L, identifying it as a critical regulator of the DNA repair process and a steward of genomic integrity.

## Introduction

Histone modifications and histone variant exchange are essential components of DNA repair pathways in eukaryotes. Phosphorylation of the histone variant H2A.X, termed γH2A.X, is an early histone modification that contributes to sensing DSB. The propagation of γH2A.X surrounding the break site leads to the recruitment of the remodeling complex containing p400/Tip60 via the scaffolding protein MDC1 (Downs et al. 2004; Xu et al. 2010). The p400-catalyzed exchange of H2A for H2A.Z into the nucleosome at DSB sites is accompanied by local histone acetylation and histone ubiquitination (Xu et al. 2012). These modifications serve as signals for the recruitment of several DSB repair factors, including BRCA1 and 53BP1, proteins necessary for both HDR and NHEJ repair.

The extent to which histone modifications contribute to histone variant exchange at DSB sites is not well understood. Histone modifications are carried out by several, often redundant, enzymes. Thus, the development of models to study epigenetic marks in isolation has been difficult. An exception to this is H3K79 methylation, which is carried out by a single enzyme called disruptor of telomeric silencing 1-like, Dot1L (Dot1 in yeast). Dot1L is the only known H3K79 methyltransferase in eukaryotic cells and is responsible for deposition of mono-, di- and tri-methyl marks (Feng et al. 2002; Ng et al. 2002). Accumulating evidence from eukaryotes suggests that H3K79 methylation may be required for maintaining genome integrity, particularly in yeast, potentially by its role in the DNA damage response (Huyen et al. 2004; Giannattasio et al. 2005; Wysocki et al. 2005; Botuyan et al. 2006; Lazzaro et al. 2008; Conde et al. 2009; FitzGerald et al. 2011; Wakeman et al. 2012). However, the precise role of H3K79 methylation in the DNA damage response, especially in higher eukaryotes, is unclear. Some studies have linked H3K79 methylation to recruitment of the critical repair factor 53BP1 to DNA double strand breaks (Huyen et al. 2004; Wakeman et al. 2012), while others have implied a lesser role for H3K79 methylation in 53BP1 recruitment (Botuyan et al. 2006). A study with chicken cells showed no role for H3K79 in 53BP1 recruitment, and the effects of H3K79 methylation on DNA repair were attributed, at least in part, to alterations in the expression of DNA damage repair factors (Huyen et al. 2004; FitzGerald et al. 2011). In this study we used *Dot1l^-/-^* mouse embryonic cells (Feng et al. 2010) and CRISPR/Cas9-directed *DOT1L* mutation in human cells to examine the role of Dot1L in DNA repair and genome stability. We find that the absence of Dot1L-catalyzed H3K79 methylation indeed results in genomic instability in mammalian cells. Further, we show that this phenomenon can be attributed to altered kinetics of both 53BP1 and BRCA1 recruitment during early DNA repair events, decreased H2A.Z exchange at sites of DSB, and a profound decrease in HDR activity, which collectively result in a striking decrease in DNA repair fidelity.

## Results

### Cells from mice lacking Dot1L accumulate DNA damage and exhibit genomic instability

The *Dot1l^-/-^* mice lack H3K79 methylation and die at E10.5, in part due to defects in hematopoiesis caused by dysregulation of an erythroid-specific transcriptional program (Feng et al. 2010). To determine whether hematopoietic progenitors derived from *Dot1l^-/-^* mice accumulate DNA damage, cells from E10.5 yolk sacs were cultured in media containing cytokines that promote growth of erythroid and myeloid progenitors: SCF, IL-3, IL-6, and EPO. After 4 days of growth, cells were analyzed for endogenous DNA damage. In an alkaline comet assay, hematopoietic progenitors from *Dot1l^-/-^* mice showed significant increases in DNA damage when compared to progenitors derived from wild-type (WT) littermates (Fig. 1a).

**Alvarez_Fig_1.**
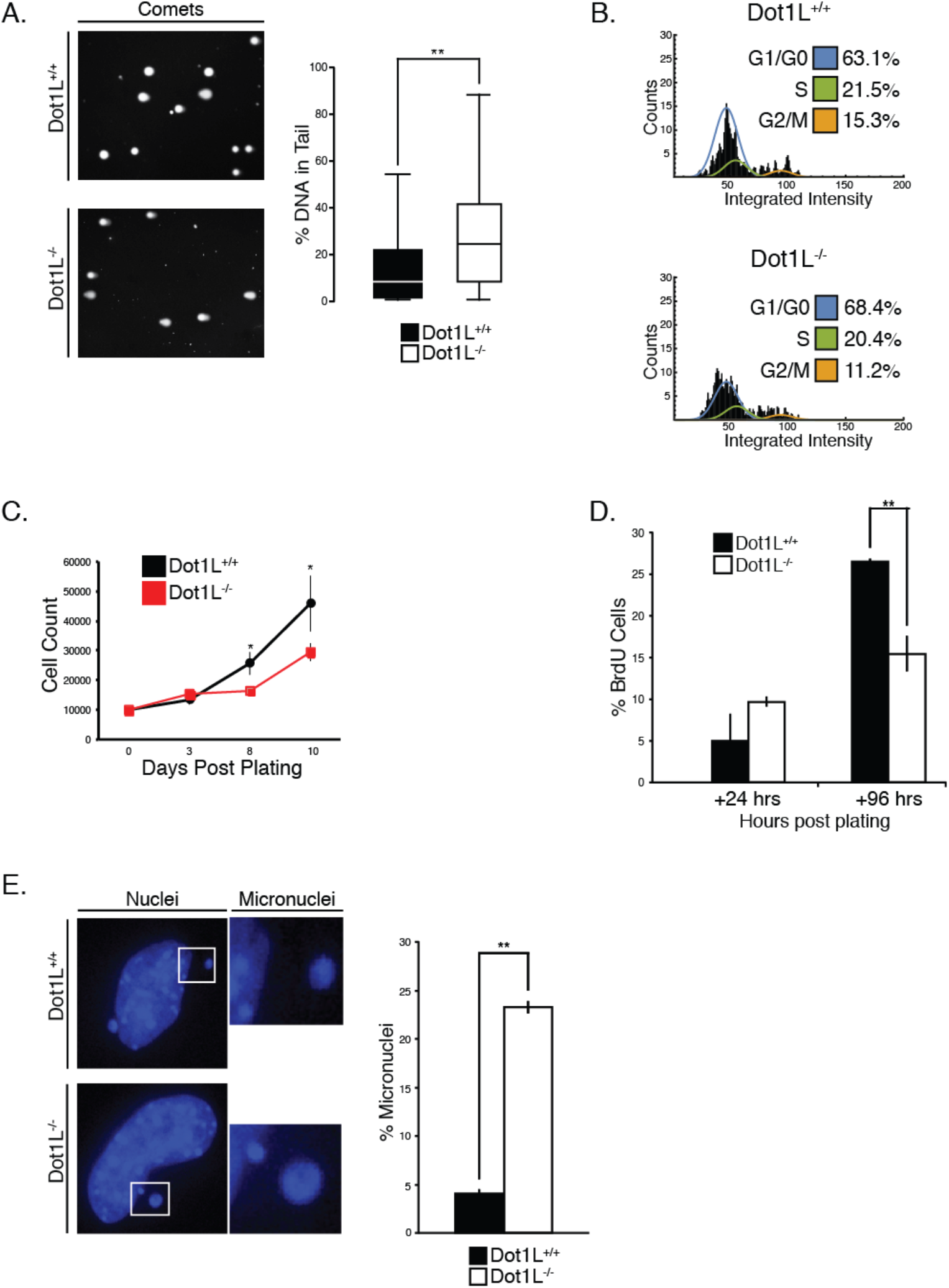
Marks of genome instability increase in Dot1L^-/-^ mouse embryonic cells. (***A***) Single cell electrophoresis (comet assay) on yolk sac cells isolated from E10.5 *Dot1L^-/-^* and wild-type mice (see *Methods*); left panel, representative fluorescence images; right panel, DNA content in the tail was quantified as described in *Methods*. Data represent the mean ± sem, n=3. (***B***) Embryonic cells from E10.5 mice of the indicated genotype were isolated and cultured for 48 hrs. Cell cycle analysis was performed as described in *Methods*. (***C***) Growth kinetics of cells isolated from the embryo proper of E10.5 mice. Data are mean ± sem, n=3. (***D***) Percentage of Hoechst 33342 stained cells positive for BrdU isolated from the embryonic cells from E10.5 mice of the indicated genotype were isolated, cultured for 2-5 days. (***E***) Hoechst 33342 stained nuclei. Left panel, representative fluorescence images from each genotype (40X magnification); right panel, micronuclei were quantified using Mathematica software. Data are mean ± sem, n=3. *\*p < 0.05, **p < 0.01*.

The *Dot1l^-/-^* embryos are smaller than wild-type littermates, suggesting that the absence of Dot1L might result in DNA damage in cell types other than hematopoietic progenitors (Feng et al. 2010). To examine this, we assessed proliferation and accumulation of DNA damage in cells isolated from the embryo proper. Somatic cells isolated from E10.5 *Dot1l^-/-^* embryos had a delay in cell cycle progression, as demonstrated using fluorescence microscopy (Roukos et al. 2015) and an image analysis pipeline we termed MANA (machine autonomous nuclei analyzer) (Suppl Fig. 1). The overall proportion of *Dot1l^-/-^* cells in G1/G0 was greater and the fraction in G2/M was lower compared to cells from wild-type controls (Fig. 1b). The G1/G0 accumulation recapitulated the results reported for *Dot1l^-/-^* yolk sac progenitors in erythroid differentiation cultures (Feng et al. 2010). The Dot1L^-/-^ cell proliferation defects were detrimental, because fewer cells were present after three days in culture (Fig. 1c). We corroborated the proliferation defects using a microscopy-based BrdU incorporation assay, in which cells isolated from E10.5 embryos were pulsed for 24 hrs. Three days after isolation, there were fewer BrdU-positive cells derived from *Dot1l^-/-^* embryos compared to cells from wild-type littermates (Fig. 1d). We also observed that cells isolated from *Dot1l^-/-^* embryos have an increased occurrence of micronuclei, a marker of genomic instability (Fig. 1e) (Crasta et al. 2012; Zhang et al. 2015). These results indicate the presence of DNA damage, genome instability, and defective proliferation in Dot1L-deficient mouse embryos, similar to that observed in the hematopoietic progenitors (Fig. 1a).

### The loss of Dot1L-catalyzed H3K79 methylation results in specific defects in high fidelity HDR

It is possible that the absence of Dot1L results in detectable DNA damage due to deleterious effects on the DNA repair process. To test this hypothesis, we examined whether cells isolated from E10.5 embryos faithfully repair DNA following genotoxic stress. After low-dose ionizing radiation (IR), cells isolated from E10.5 *Dot1l^-/-^* embryos and wild-type littermates have similar amounts of DNA damage at 24 h post IR (Fig. 2a). However, at 72 and 96 h post-IR, cells from *Dot1l^-/-^* embryos have more accumulated DNA damage than wild-type cells (Fig. 2a). At 96 h after IR treatment, both *Dot1l^-/-^* and wild-type cells continue to proliferate, albeit at a slower rate compared to untreated controls (Fig. 2b). This observation suggests that cells from E10.5 *Dot1l^-/-^* embryos continue to proliferate despite having DNA damage, which may account for the increased levels of endogenous DNA damage that we observe in *ex vivo* culture of embryonic tissue (Fig. 1a,d).

**Alvarez_Fig_2.**
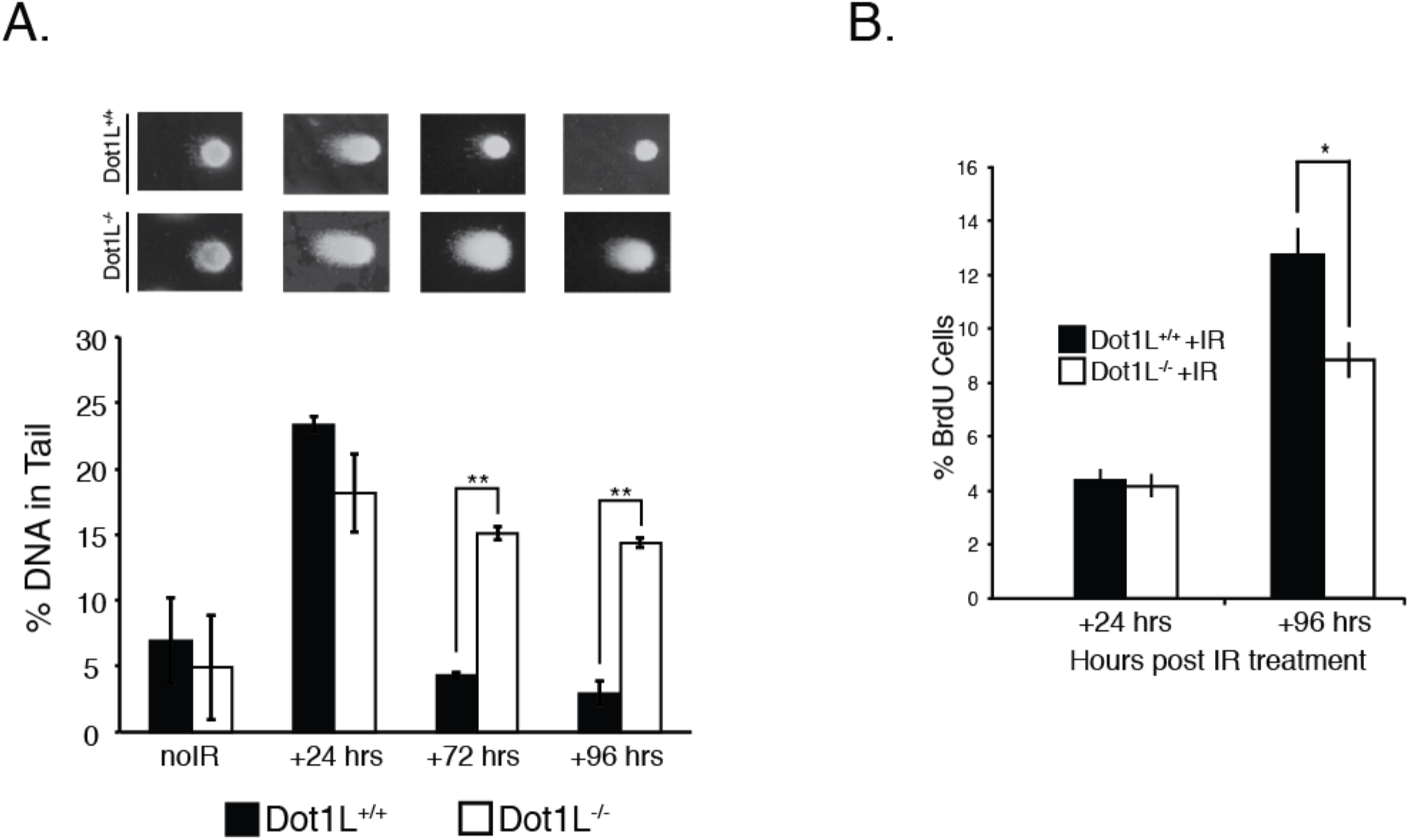
DNA damage repair capacity is reduced in Dot1L^-/-^ mouse embryonic cells. (***A***) Mean of % DNA tail content from a comet assay of E10.5 embryonic cells exposed to 2 Gy of IR. Data are mean ± sem, n=3. *\*p < 0.05, **p < 0.01*. (***B***) BrdU assay of cells isolated from the embryo proper of E10.5 mice. Data are mean ± sem, n=3. *\*p < 0.05, **p < 0.01*. (***C***) Mean % of BrdU positive cells from E10.5 embryonic cells exposed to 2 Gy of IR. Data are mean ± sem, n=3.

The accumulated DNA damage in the absence of Dot1L supports the idea that Dot1L-deficient cells have defective DNA repair. To examine this phenomenon further, we measured IR-induced γH2A.X foci, which mark DNA damage and are required for proper spatial and temporal recruitment of DDR factors. In *Dot1l^-/-^* cells exposed to 2 Gy of IR, more γH2A.X DDR foci were present at all time points measured compared to wild-type cells, and the number of foci increased over 24 h in the mutant cells (Fig. 3a). These data indicate that *Dot1l^-/-^* cells are capable of detecting DNA damage, but the damage is not being repaired.

**Alvarez_Fig_3.**
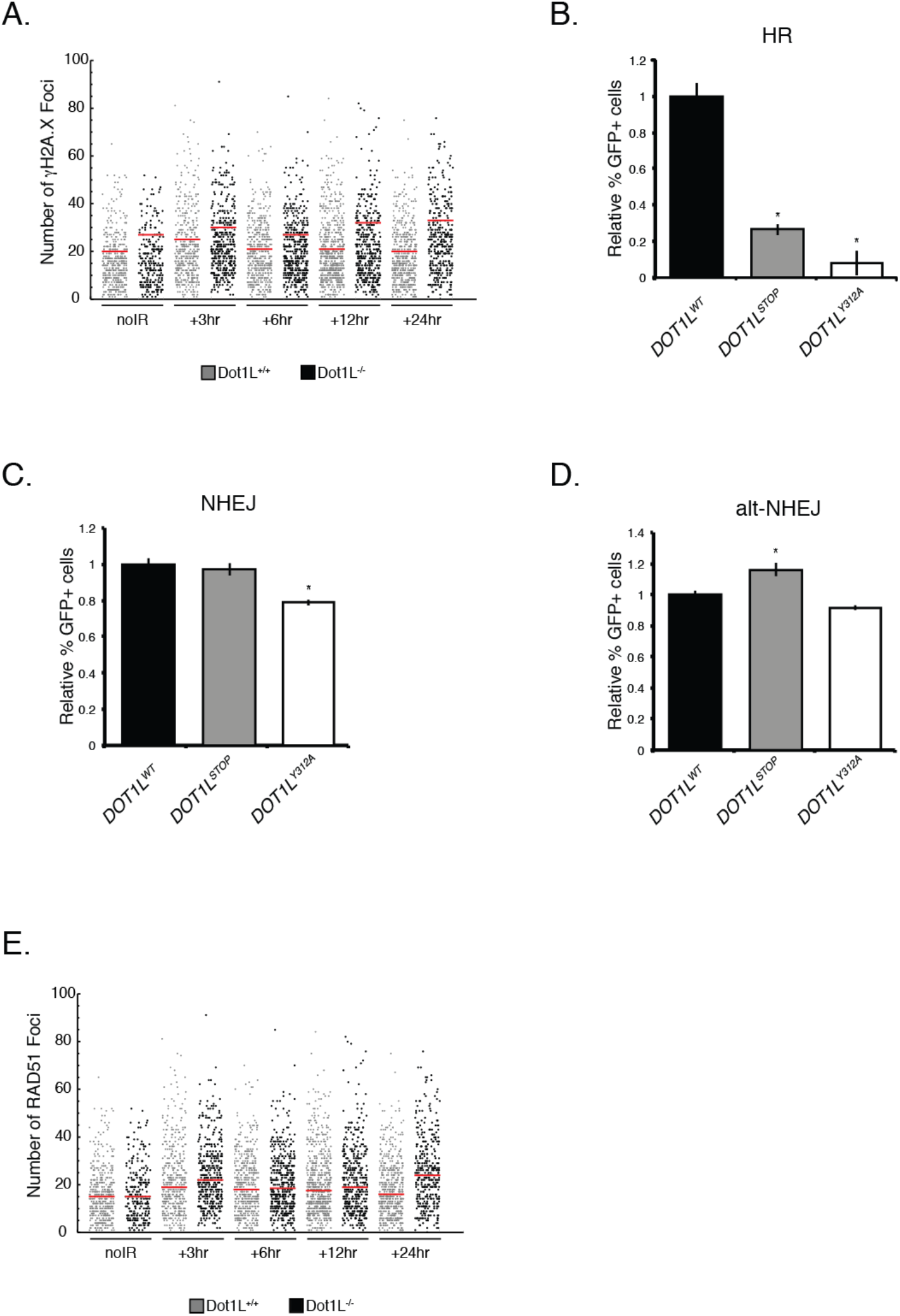
H3K79 methylation deficient cells have altered DNA damage repair pathway usage. (***A***) Distribution of γH2A.X foci in E10.5 embryonic cells, with either no IR exposure (noIR), or at 3, 6, 12, and 24 hrs after exposure to 2 Gy of IR. Horizontal bars represent median numbers of foci per cell. (***B***) Stable cell lines expressing the DR-GFP reporter transfected with the I-Sce1 endonuclease. Data are mean ± sem, n=3. (***C***) Stable cell lines expressing the E5J-GFP reporter transfected with the I-Sce1 endonuclease. Data are mean ± sem, n=3. (***D***) Stable cell lines expressing the E2J-GFP reporter transfected with the I-Sce1 endonuclease. Data are mean ± sem, n=3. *\*p < 0.05*. (***E***) Distribution of RAD51 foci in E10.5 embryonic cells, with either no IR exposure (noIR), or at 3, 6, 12, and 24 hrs after exposure to 2 Gy of IR. Horizontal bars represent median numbers of foci per cell.

To measure DNA repair in Dot1L-deficient cells directly, we used three different reporter systems to assess effects on the three major repair pathways: DR-GFP for HDR, E5J-GFP for NHEJ, and E2J-GFP for alt-NHEJ (Pierce et al. 1999; Bennardo et al. 2008). For these experiments we used CRISPR/Cas9 (Jinek et al. 2012; Mali et al. 2013) to generate both cells that lack expression of Dot1L protein (*DOT1L^Stop^*) and cells that express a mutant, non-functional form of Dot1L (*DOT1L^Y312A^*) (Suppl Fig. 2). The latter mutation, which is known to inactivate Dot1L methyltransferase activity (Min et al. 2003), facilitates evaluation of any H3K79 methylation-independent function of the Dot1L protein (e.g., its capacity to interact with other proteins). As expected, both *DOT1L^Stop^* and *DOT1L^Y312A^* HEK293T cells have decreased H3K79 mono- and di-methylation marks, while only *DOT1L^Stop^* reduces DOT1L protein expression (Suppl Fig. 2h). The residual low level of Dot1L protein in the *DOT1L^Stop^* cells is likely the result of small amounts of long-lived protein, since sequencing confirmed insertion of stop codons in all three reading frames in both alleles (not shown).

Both loss of Dot1L (*DOT1L^Stop^*) and loss of Dot1L methyltransferase activity (*DOT1L^Y312A^*) resulted in a dramatic reduction in HDR activity (Fig 3b). Notably, the DNA repair defects were highly specific for the high fidelity HDR pathway, as activities of both the NHEJ and alt-NHEJ repair pathways were maintained in *DOT1L^Stop^* and *DOT1L^Y312A^* cells (Fig. 3c-d). These data indicate that in the absence of H3K79 methylation activity, cells preferentially utilize more error-prone DNA repair pathways (NHEJ and alt-NHEJ) to repair DSB rather than the HDR pathway. The absence of high fidelity DNA repair would be predicted to result in genomic instability (McVey et al. 2016), which is precisely the phenotype exhibited by cells with loss of Dot1L (Fig 1).

### Dot1L-catalyzed H3K79 methylation affects timing of assembly of DNA repair foci

To elucidate the underlying cause of the DNA repair defect, we examined expression of DNA damage response (DDR) genes. Dot1L-catalyzed H3K79 methylation is associated with transcriptional activation, and prior reports had suggested that transcriptional regulation is a potential means by which Dot1L could influence DNA repair (Mohan et al. 2010; FitzGerald et al. 2011). Surprisingly, however, gene expression analysis for DDR pathway factors in cells isolated from E10.5 *Dot1l^-/-^* embryos demonstrated *elevated* expression of multiple factors involved in both sensing and repairing DNA damage (Fig. 4a-c). Thus, defective DNA repair in the absence of Dot1L seems unlikely to be primarily the result of diminished expression of relevant repair factors.

**Alvarez_Fig_4.**
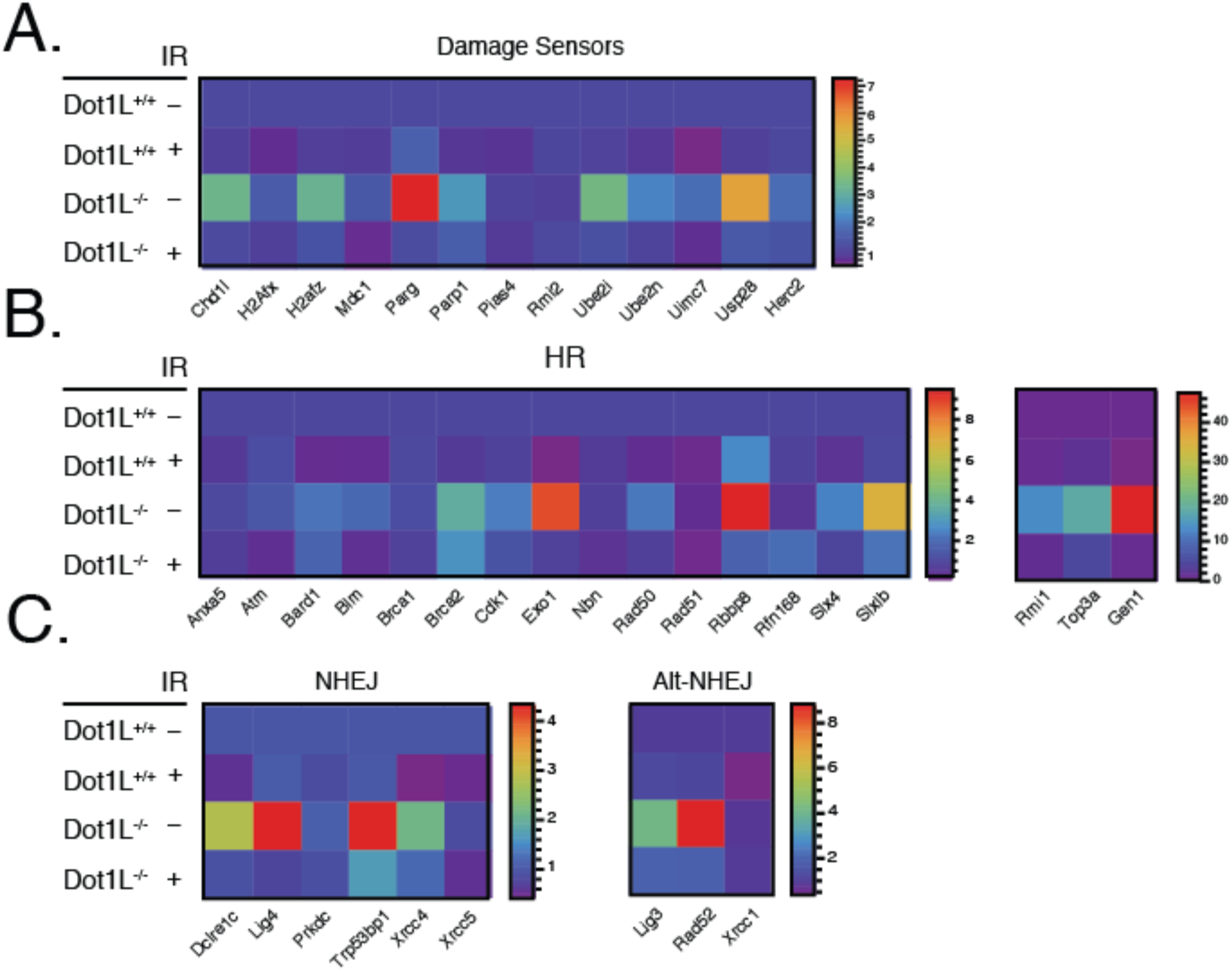
Expression of DNA damage response factors is elevated in Dot1L^-/-^ mouse embryonic cells. Expression of DNA damage sensors (***A***), HR factors (***B***), and NHEJ factors (***C***) from E10.5 embryonic cells exposed to 2 Gy of IR. Scale indicates the ΔCt value.

The timing of the assembly of pathway-specific, DNA damage repair factors at DSB sites can influence DNA repair pathway utilization (Luo et al. 2012; Chapman et al. 2013; Tang et al. 2013; Watanabe et al. 2013). So, we used immunofluorescence microscopy to assess the effects of Dot1L loss on the loading of DDR factors. Since Dot1L-deficient cells displayed a HDR-specific defect, we examined IR-induced foci of RAD51, a protein that reportedly coordinates HDR activity (Thacker 2005; Shrivastav et al. 2008; Lisby and Rothstein 2009; Shrivastav et al. 2009). Somewhat surprisingly, in Dot1L^-/-^ cells exposed to 2 Gy of IR, RAD51 foci were found to be slightly elevated when compared to wild-type cells at 24 h post irradiation using a Mann-Whitney test (Fig. 3e).

We next focused on the kinetics of 53BP1 and BRCA1 DDR foci formation, since these factors are known to be central to the choice of HDR versus NHEJ for DNA repair (Daley and Sung 2014; Panier and Boulton 2014; Ochs et al. 2016). Also, H3K79 methylation has been reported to influence 53BP1 targeting to DSB (Huyen et al. 2004). We treated wild-type, *DOT1L^Stop^*, and *DOT1L^Y312A^* cells with IR and analyzed 53BP1 and BRCA1 foci formation at different time points (Fig. 5a,b). At 30 and 60 min post IR, *DOT1L^Stop^* and *DOT1L^Y312A^* cells have reduced 53BP1 foci compared to wild-type HEK293T cells, but by 90 min have similar numbers of foci per cell (Fig. 5a). BRCA1 foci levels for *DOT1L^Stop^* and wild-type cells are similar at all time points, however, *DOT1L^Y312A^* cells have a significant decrease in foci abundance at 60 min post IR (Fig. 5b). We compared the co-localization of 53BP1 and BRCA1 following IR treatment. We observed reduced 53BP1/BRCA1 co-localization events in both the *DOT1L^Stop^* and *DOT1L^Y312A^* mutant cells 30 min post IR (Fig. 5c). It has been reported that 53BP1 limits the end resection activity of BRCA1 during the S phase of the cell cycle (Chapman et al. 2013; Ochs et al. 2016). We determined whether the ratios of 53BP1 to BRCA1 foci were differentially affected in *DOT1L^Stop^* and *DOT1L^Y312A^* cells following exposure to IR (Fig. 5d). At 30 min post IR, both *DOT1L^Stop^* and *DOT1L^Y312A^* cells display reduced ratios of 53BP1/BRCA1 during S phase (Fig. 5d). We also observed an increase in the population of BRCA1-positive S phase cells 30 min post IR in both mutant cell lines, but no compensatory increase in 53BP1positive S phase cells (Fig. 5e, f). We did not observe this phenomenon in the G1 populations of either cell line (Suppl Fig. 3). The decrease in co-localization of 53BP1/BRCA1 and increased amounts of BRCA1 positive S-phase cells indicate that the coordination between 53BP1 and BRCA1 in fine-tuning the DSB DDR is reduced in the absence of H3K79 methylation.

**Alvarez_Fig_5.**
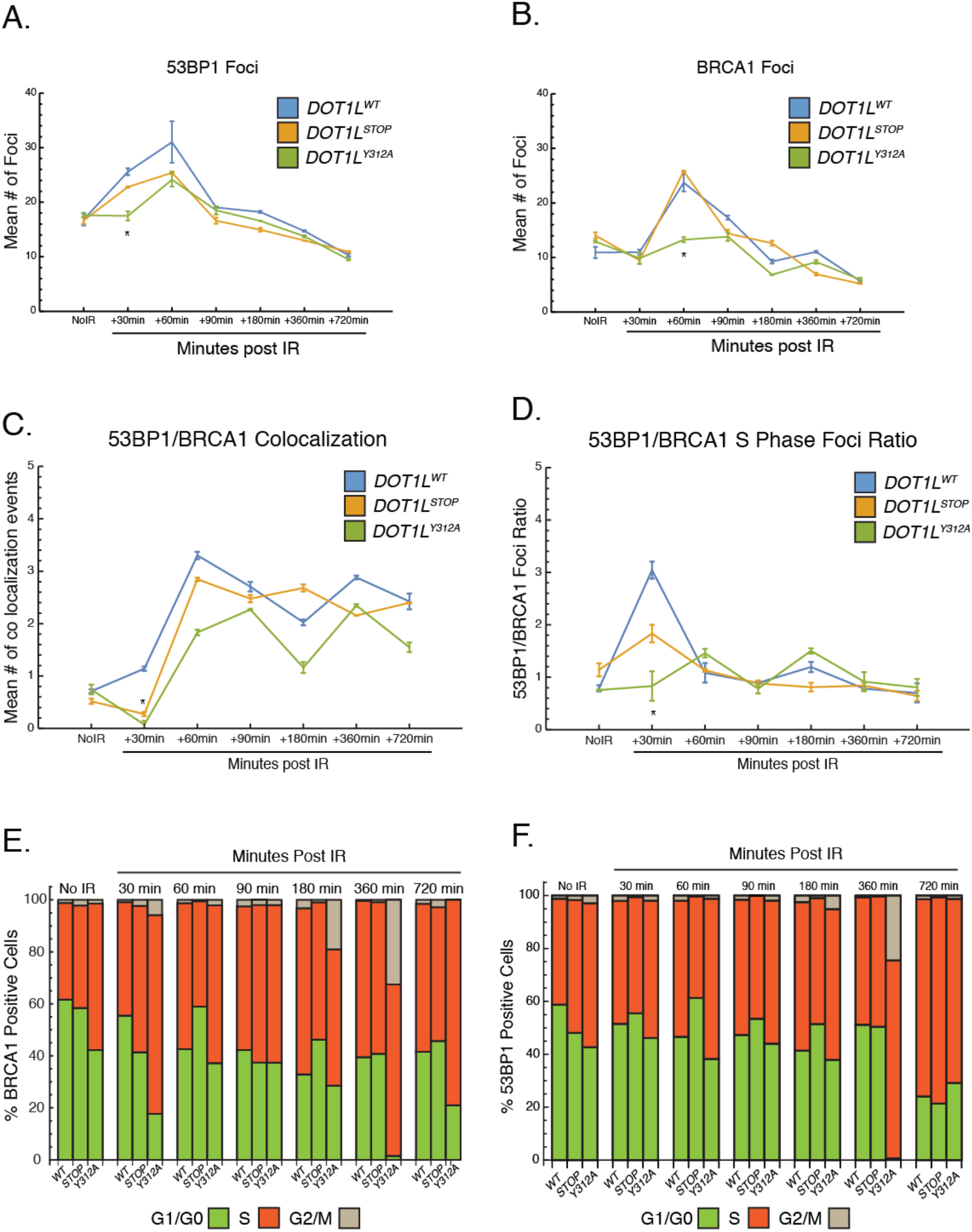
H3K79 methylation deficient cells have altered 53BP1/BRCA1 foci kinetics. Mean numbers of 53BP1 (***A***) and BRCA1 (***B***) foci in cells exposed to 10 Gy of IR. Data represent mean ± sem, n=3. *\*p<0.05* wild-type vs. *DOT1L^Y312A^*. Mean of 53BP1/BRCA1 co-localized foci (***C***) in cells exposed to 10 Gy of IR. Data are mean ± sem, n=3. *\*p<0.05* wild-type vs. *DOT1L^STOP^*, wild-type vs. *DOT1L^Y312A^*. Mean of the ratio of 53BP1/BRCA1 foci (***D***) in cells in the S phase of the cell cycle exposed to 10 Gy of IR. Percentage of BRCA1 (***E***) and 53BP1 (***F***) positive cells in each phase of the cell cycle following exposure to 10 Gy of IR.

### H3K79 methylation is required for efficient H2A.Z incorporation into nucleosomes at sites of DSB

To further clarify the mechanism by which Dot1L-catalyzed H3K79 methylation influences HDR, we also examined histone changes at DSB. To accomplish this, we developed a technique that allows evaluation of histone dynamics at DSB without prior knowledge of the sequence surrounding the DSB or overexpression of endonucleases: broken-end ligation chromatin immunoprecipitation (BEL-ChIP). This technique facilitates identification of proteins specifically enriched at DSB (Suppl Fig. 4a). Using BEL-ChIP, we first assessed H4 acetylation at DSB. Evidence suggests that H4 acetylation can alter the relative ratios of BRCA1 and 53BP1 at DSB sites by preventing the association of 53BP1 (Tang et al. 2013). Moreover, Dot1L-dependent H3K79 methylation has been reported to facilitate H4 acetylation (Gilan et al.). In the absence of IR, similar H4 acetylation was present in chromatin. After IR, there was a mild decrease in global H4 acetylation in both *DOT1L^Stop^* and *DOT1L^Y312A^* cells that was most evident after 60 min (Suppl Fig 4b). However, BEL-ChIP revealed no significant difference in H4 acetylation at DSB between wild-type, *DOT1L^Stop^* and *DOT1L^Y312A^* cells (Suppl Fig. 4c). Thus, it is not likely that changes in H4 acetylation at DSB account for the defective utilization of the HDR pathway.

We next examined histone H2A.Z variant exchange, since recent work has indicated that exchange of this protein into the nucleosomes surrounding DSB sites can influence DNA repair pathway utilization (Xu et al. 2012). We first assessed the effects of H3K79 methylation on the stability of H2A.Z-containing nucleosomes, since H3K79 methylation has been reported to affect nucleosome structure (Lu et al. 2008). Without IR, similar amounts of H2A.Z are found tightly associated with chromatin, regardless of Dot1L status (Fig. 6a). However, after IR, H2A.Z is more stably associated with DNA after 1 and 2 hours in both *DOT1L^Stop^* and *DOT1L^Y312A^* cells (Fig. 6a).

**Alvarez_Fig_6.**
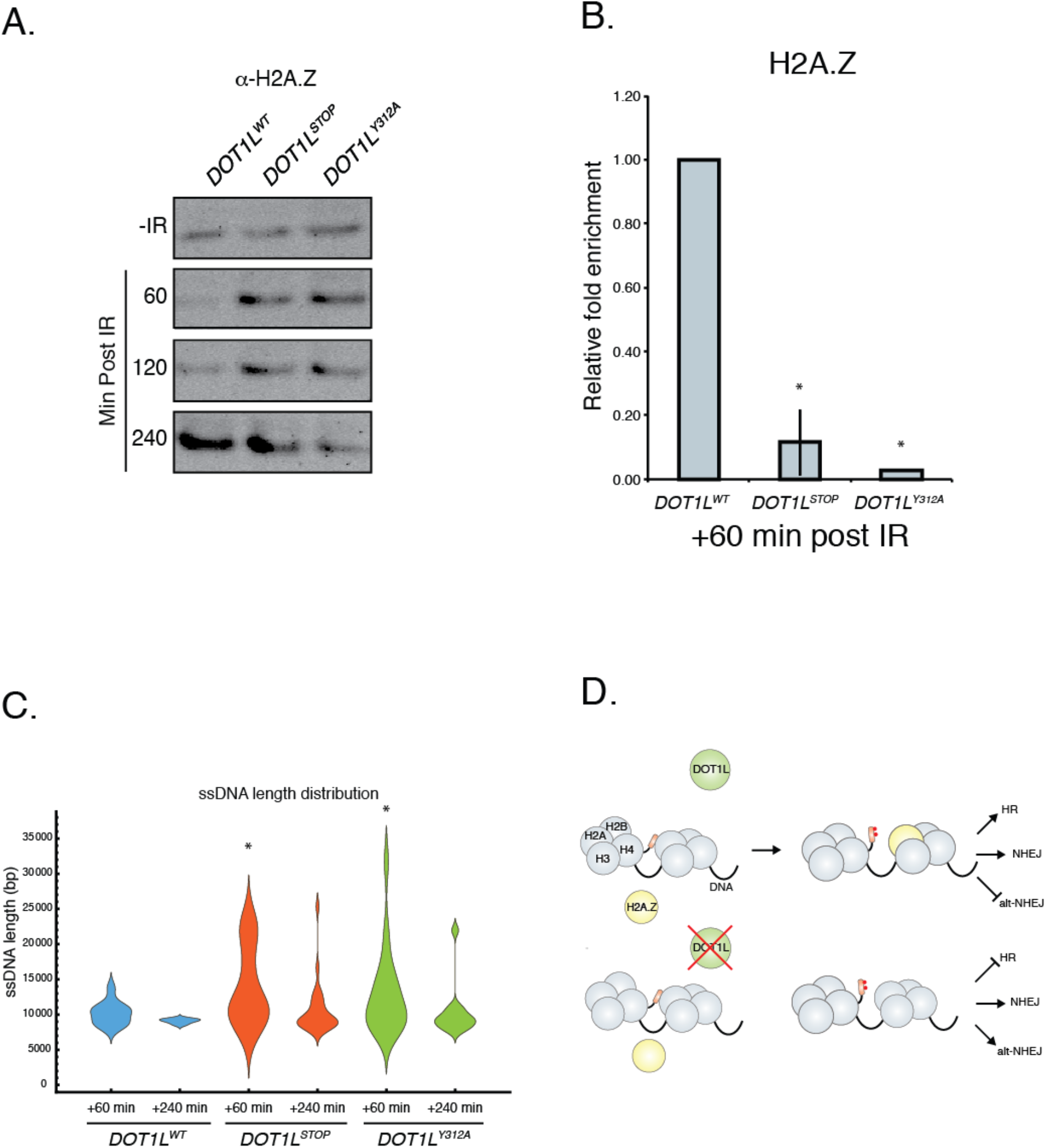
H3K79 methylation deficient cells have defects in H2A.Z incorporation at sites of double strand breaks. (***A***) Histone stability assay following exposure to 10 Gy of IR. Western blot of histone H2A.Z. (***B***) BEL-ChIP of histone H2A.Z at sites of double strand breaks. Data are mean ± sem, n=3. (***C***) Length distribution of ssDNA after exposure to 10 Gy IR. Calculated lengths were normalized to No IR input for each sample (***D***) Model for the role of DOT1L in DSB repair.

The tight association of H2A.Z in the absence of H3K79 methylation may prevent H2A.Z mobilization and efficient exchange at DSB. To test this hypothesis, we used BEL-ChIP to quantify H2A.Z at DSB. This analysis demonstrated that both *DOT1L^Stop^* and *DOT1L^Y312A^* cells have markedly decreased enrichment of H2A.Z at DSB sites following exposure to IR (Fig. 6b). These data suggest that efficient H2A.Z exchange into nucleosomes at DSB sites requires H3K79 methylation and may be related to altered stability and mobilization of H2A.Z.

Another consequence of H2A.Z nucleosome incorporation at DSB is restriction of the activity of the end resection machinery, which facilitates the utilization of HDR (Xu et al. 2012). In the absence of H2A.Z, end resection is thought to be over-active, favoring the generation of single stranded DNA (ssDNA), which is a poor substrate for HDR (Xu et al. 2012). To measure the length of ssDNA surrounding DSB, we developed an assay using Oxford Nanopore technology (Suppl Fig 4d). The length of ssDNA was assessed in WT and Dot1L mutant cells after IR. This analysis revealed that both *DOT1L^Stop^* and *DOT1L^Y312A^* mutants generated significantly longer ssDNA than wild-type cells after IR (Fig. 6c). This elongated ssDNA would limit utilization of HDR, which is consistent with the observation that both *DOT1L^Stop^* and *DOT1L^Y312A^* cells have defective HDR activity (Fig 3).

## Discussion

In this study we confirm a role for Dot1L-dependent H3K79 methylation in DNA repair and extend previous findings in yeast, demonstrating a specific role for Dot1L in HDR and maintenance of genomic stability in mammalian cells. Dot1L is associated with transcriptional activation (Steger et al. 2008; Mohan et al. 2010), an activity that has been postulated to contribute to effects on DNA repair. However, we demonstrate that the mechanism underlying the influence of Dot1L on DNA repair and genome stability involves regulation of H2A.Z variant exchange at DSB: the lack of H2A.Z at DSB allows enhanced end resection activity and promotes the generation of long ssDNA, which can influence DNA repair pathway utilization and decrease HDR (Xu et al. 2012; Gursoy-Yuzugullu et al. 2015). In the absence of H3K79 methylation, diminished HDR and unimpeded utilization of the error-prone NHEJ and alt-NHEJ pathways contributes to genomic instability (Fig. 6d). Also, the absence of Dot1L may affect the DNA damage checkpoint, similar to that described in yeast (Giannattasio et al. 2005; Wysocki et al. 2005). Cells from *Dot1l^-/-^* embryos continue to proliferate despite having DNA damage (Fig 1a,d), which may also contribute to overall genomic instability.

The precise mechanism by which H3K79 methylation facilitates H2A.Z exchange is not certain, but our data suggest that it may be related to effects on nucleosome stability. In the absence of H3K79 methylation, H2A.Z displayed increased stability in nucleosomes from cells exposed to IR (Fig 6a), which could prevent mobilization for exchange after DSB. This conclusion is consistent with data showing that H3K79 methylation alters nucleosome surface properties and structure (Lu et al. 2008).

In addition to direct structural effects of H3K79 methylation, Dot1L may also influence H2A.Z nucleosome stability by affecting nucleosome composition. Evidence suggests that H3K79 methylation occurs predominantly on H3.3 (McKittrick et al. 2004), so it possible that H3K79 methylation facilitates H3.3/H2A.Z assembly. Since H3.1/H2A.Z containing nucleosomes have been shown to be as stable as H3.1/H2A containing nucleosomes (Jin and Felsenfeld 2007), the net effect of the absence of H3K79 methylation could be a decrease in the H3.3/H2A.Z nucleosomes and an increase in the stable H3.1/H2A.Z nucleosomes, which could result in reduced mobilization of H2A.Z.

Our findings showing altered H2A.Z incorporation into DSB are different from recent studies analyzing H2A.Z in some respects (Gursoy-Yuzugullu et al. 2015). In that previous study, the authors showed only a transient increase in H2A.Z in foci after DNA damage, while we have observed H2A.Z enrichment at DSB at 60 min (Fig 6b). This difference may be related to the methodology used to measure H2A.Z exchange. In the prior paper, H2A.Z association was measured by immunofluorescence microscopy, while in this study we developed a method (BEL-ChIP) that measures enrichment of H2A.Z specifically at DSB. Another difference in our studies is the method of DSB induction: we used whole cell irradiation and the other study used laser microirradiation.

Cells with deficient H3K79 methylation also show altered DNA repair foci formation after IR. Foci with RAD51 were slightly elevated in *Dot1l^-/-^* cells. This could be the result of increased ssDNA substrate present in the *Dot1l^-/-^* cells. RAD51 binds to ssDNA, forming filaments necessary for HDR to take place (Liu et al. 2010). However, as the length of ssDNA increases, more mutagenic repair pathways are used (Richardson et al. 2004; Ochs et al. 2016). Dot1L-deficient cells also show altered kinetics of 53BP1 and BRCA1 repair foci formation. BRCA1 and 53BP1 are recruited to DSB prior to RAD51 and define the region surrounding the DSB. Alterations of these factors have been associated with increased ssDNA and the utilization of highly mutagenic DNA repair pathways (Xu et al. 2012; Chapman et al. 2013; Ochs et al. 2016).

H3K79 methylation correlates with active gene transcription (Steger et al. 2008; Mohan et al. 2010). Similarly, H2A.Z is enriched at the promoters of actively transcribed genes (Subramanian et al. 2015). It is possible that the co-localization of H3K79 methylation and H2A.Z mark regions of the genome that require HDR. Rearrangements in MLL genomes, which result in the redistribution of H3K79 methylation, have normal efficiencies of NHEJ, HDR, and alt-NHEJ (Castano et al. 2016). This would suggest that H3K79 methylation is an epigenetic mark required for histone variant exchange and HDR in a variety of contexts. Supporting this hypothesis, we have observed that inhibiting H3K79 methylation in MCF7 cells results in apoptosis (data not shown). Determining which epigenetic marks facilitate histone variant exchange during DNA repair processes could provide insight into the formulation of novel therapeutic combinations to treat malignancies beyond leukemia.

## Methods

### Dot1L knockout mice

Dot1L^-/-^ mice were previously described (Feng et al. 2010).

### *Ex vivo* erythroid differentiation assay

Cells from E10.5 *Dot1l^-/-^* and wild-type yolk sacs were cultured in M3434 methylcellulose medium (StemCell Technologies) containing the cytokines SCF, IL-3, IL-6, and EPO, which promote the growth of erythroid and myeloid progenitors.

### Cell Culture

Wild type HEK293T and mutant HEK293T *DOT1L^STOP^* and *DOT1L^Y312A^* cell lines were maintained in 10% FBS, DMEM, Pen/Strep in 37°C, 5% CO_2_. Embryonic day 10.5 (E10.5) embryos were obtained from timed matings between heterozygous Dot1L mutant mice, cells were isolated from the whole embryo and cultured in DMEM, high glucose, L-glutamate, Sodium Pyruvate (Life Technologies, 11995-040), 10% fetal bovine serum (Life Technologies 10082-147), supplemented with 20 mM HEPES (Life Technologies 15630-080), L-glutamate, and essential amino acids.

### Cas9 Targeting of HEK293T cells

*DOT1L^Stop^* and *DOT1L^Y312A^* cell lines were generated by Cas9/ssDNA directed cleavage/repair. The CRISPR/Cas9 vector pX330-U6-Chimeric_BB-CBh-hSpCas9 was a gift from Feng Zhang (Addgene plasmid # 42230) (Cong et al. 2013). The CRISPR/Cas9 vector was modified to express an eGFP-P2A-Cas9 fusion gene using SLiCE (Seamless Ligation Cloning Extract) method (Zhang et al. 2012). Briefly, overlapping PCR was used to generate eGFP-P2A cDNA with arms of homology for pX330 promoter and N-terminus of hSpCas9. 2 μL of SLiCE was mixed with eGFP-P2A cDNA and digested pX330 in a 1:4 molar ratio and incubated at 37°C for 1 hr. 2 μL of the reaction was transformed into *E. coli.* Single clones were isolated and verified by Sanger sequencing. SLiCE strain of *E. coli* was a gift from Yongwei Zhang (Albert Einstein College of Medicine). The guide RNA targeting the *DOT1L* locus (sequence below) was cloned into eGFP-P2A-Cas9 using BbsI. The same guide RNA was used to generate both *DOT1L^Stop^* and *DOT1L^Y312A^* cell lines. DOT1L Y312 was chosen for targeting because previous work demonstrated that mutation to this residue disrupted H3K79 methyltransferase activity, and it satisfied CRISPR/Cas9 targeting parameters (Min et al. 2003; Hsu et al. 2013). A 150 nt single stranded oligonucleotide, antisense to transcription, was used to direct repair of the Cas9 cleaved locus by destroying the PAM, adding a DdeI restriction site and either adding three tandem stop codons (*DOT1L^Stop^*) or nucleotides converting Tyrosine 312 to Alanine (*DOT1L^Y312A^*). HEK293T (ATCC) cells were transfected with a 1:1 molar ratio of eGFP-P2A-Cas9 and single stranded oligonucleotide using Effectine (Qiagen) in a six-well dish. Cells were sorted for eGFP expression 48 h after transfection (FACSaria II, BD). Colonies were picked 5 days later, genomic DNA was extracted, and a PCR/restriction digest with DdeI was used to initially identify DOT1L and Y312A positive clones. Western blot and Sanger sequencing was used to verify clones.

The following primers were used to PCR screen for mutations:

DOT1L Screen Forward: GTG TGG GAG AAG AGG GAA GA

DOT1L Screen Reverse: AGC AGT ACT GAG GGG CCT TG

The following primers were used for cloning the guide RNA into the CRIPSR/Cas9 vector:

DOT1L Guide Forward: CAC CGT CGA TAG TGT GCA GGT AGT

DOT1L Guide Reverse: AAA CAC TAC CTG CAC ACT ATC GAC

The following sequences are the repair templates used to generate the *DOT1L* mutants:

DOT1L Stop Repair Template:

CAG GAC GGC CCT GCG GCT GAG GCG CAG CGA GAT ACT CAC TAT GGT GCG GTC GAT AGT GTG CTA CTA CTA AGA GAC TGG CTT CCC CGT CCA CGA CAC TGA TCC TTT CAG GGG CGA GAG CTC CAC CAC GCG CAT GAT GGT GCC GAT GTC TGC

DOT1L Y312A Repair Template:

CAG GAC GGC CCT GCG GCT GAG GCG CAG CGA GAT ACT CAC TAT GGT GCG GTC GAT AGT GTG CAG GTA GGC TGA GAC TGG CTT CCC CGT CCA CGA CAC TGA GCC CTT CAG GGG CGA GAG CTC CAC CAC GCG CAT GAT GGT GCC GAT GTC TGC

### Focus formation

HEK293T *DOT1L^STOP^*and *DOT1L^Y312A^* cell lines and cells isolated from E10.5 *Dot1l^-/-^* and wild-type embryos were plated onto cover slips treated with L-Poly lysine (75 000 -150 000 MW). Cells were exposed to 10 Gy of ionizing radiation. Cells were fixed at different time points post irradiation with 4% Paraformaldehyde for 30 min at room temperature, cells were washed with 200 mM Glycine/1X Dulbecco’s phosphate-buffered saline (DPBS) for 10 min, permeabilized with 0.2 % TritonX100/1X DPBS for 30 min, blocked with 1% BSA/1X DPBS for 60 min. Mouse DDR foci were visualized by staining with γH2A.X (Abcam #ab2893,1:1000), 53BP1(Abcam #ab36823, 1:500) for 1 hr, followed by secondary antibody conjugated to Alexa Fluor 488 (ThermoFisher # A-11008, 1:1000) for 2 hrs. HEK293T cells costained for 53BP1(Abcam #ab36823, 1:500) and BRCA1 (Abcam #ab16781, 1:2000) were visualized by staining for 1 hr, followed by appropriate secondary antibodies conjugated to Alexa Fluor 488 (ThermoFisher # A-11008, 1:1000) and Alexa Fluor 594 (ThermoFisher # A-11001, 1:2500) for 2 hrs. Cells were washed with 1X DPBS/Hoechst (1μg/mL) 3 times for 5 min.

### Western antibodies

The following antibodies were used for Western blots: di-methyl H3K79 (1:1000; Abcam #ab3594), mono-methyl H3K79 (1:1000; Abcam #ab2886), histone H3 (1:1000; Abcam #ab24834), Dot1L (1:1000; Abcam #ab57827), H2A.Z (1:1000; CellSignaling #2718).

### DNA damage pathway gene expression

Total RNA was isolated from cells of E10.5 *Dot1L^-/-^* and wildtype mice that were 6 hours post ionizing radiation treatment (2 Gy). qPCR was performed using Power SYBR® Green PCR Master Mix (#4367659, Thermofisher Scientific), primers with annealing temperature of 61°C and 40 cycles on ABI 7900 system. Primers (listed below) for mouse DNA damage pathway genes were designed using IDT PrimerQuest Tool.

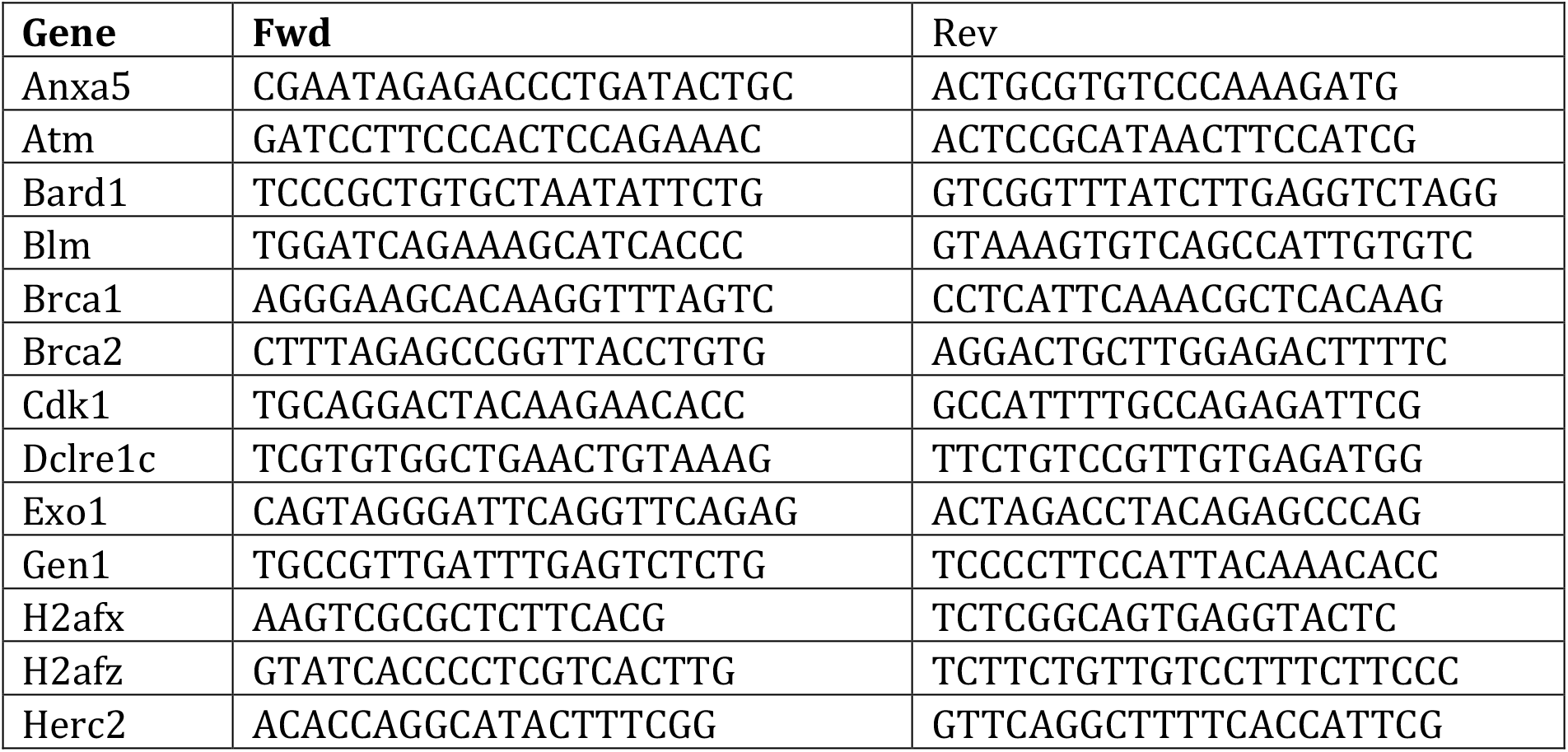
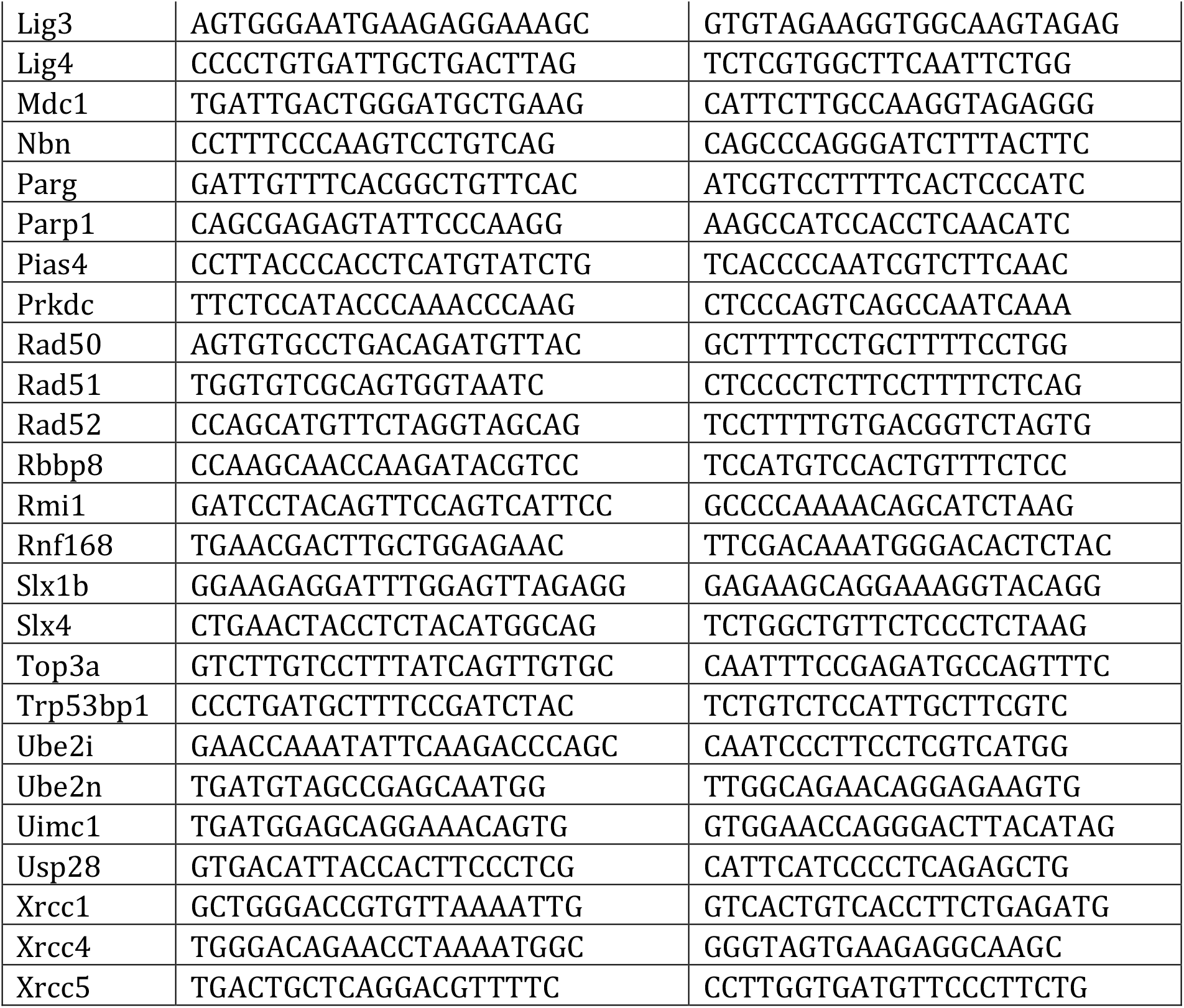

### ssDNA measurement and MinlON

Wild type HEK293T and mutant HEK293T *DOT1L^STOP^* and *DOT1L^Y312A^* cell lines were exposed to 10 Gy of ionizing radiation and total genomic DNA was isolated at 60 and 240 min after radiation exposure by adding resuspended cell pellets in 50 μL of Extraction buffer (Tris-HCl, pH 9.5, 0.1M, KCl 0.25 M, EDTA 0.01 M), 10 μL Proteinase K (20 mg/mL), 10 μL RNAseA (10 mg/mL, Fermentas). The samples were incubated at 55°C for 15 min and ethanol precipitated. The extracted DNA was digested with RNAse A and dsDNAse (ThermoFisher) overnight at room temperature. The samples were ethanol precipitated twice. ssDNA was converted to dsDNA using the following reaction 5 μL dNTP, 2μL AMP Ligase (ThermoFisher), 2μL T4 DNA polymerase, 10μL Random hexamer, 5 μL Ligase buffer. Reaction was incubated at 25°C for 10 min and 16°C for 16 hrs. The samples were precipitated and the sequencing library was prepared as described in the MinION documentation (https://nanoporetech.com/products/minion). Base calling was performed by Metrichor 2.42.2, and sequence analysis was performed using the Mathematica programming language. DNA molecules present in the samples without IR were subtracted from the +60 minute and +240 minute post IR samples, using a custom search string search algorithm that scans and removes identical sequences from fastq files.

### BEL-ChIP

The broken end linker and anchor linker (IDT) were annealed using T4 DNA Ligase Reaction Buffer (#B0202S, New England Biolabs) to a final concentration of 10 μM. The reaction was heated to 98°C for 5 min and allowed to cool to room temperature at 1 °C per minute. Wild type HEK293T and mutant HEK293T *DOT1L^STOP^* and *DOT1L^Y312A^* cell lines were plated into 10 cm dishes two days prior to treatment with ionizing radiation. On the day of experiment, cells were trypsinized, neutralized with 500 μL 10% FBS-1X DMEM, and pelleted at 300 x g for 7 min. Cells were treated with 10 Gy ionizing radiation. Cells were gently resuspended and placed in a 37°C, 5% CO_2_ incubator for 60 min. Fifty-four μL of 37% Formaldehyde (2% Final conc.) was added to the sample, and incubated with constant rotation at room temperature for 30 min. Then 62.5 μL of 2 M Glycine (125 mM final) was added to the samples and incubated with constant rotation at room temperature for 5 min. Cells were pelleted at 300 x g for 7 min at 4 °C and then washed three times with ice cold PBS (no Ca^2+^, Mg^2+^). Cells were resuspended in 1 mL of ice cold buffer A (SimpleChIP Kit #9003, Cell Signaling Technology) + 0.5 μL 1M DTT + 5 μL 200X PIC (SimpleChIP Kit #9003, Cell Signaling Technology) and incubated on ice 10 min with mixing every 3 min. Cells were harvested at 300 x g for 5 min at 4°C. The pellet was washed 3X with 500 μL NEBuffer 2.1 (#B7202S, New England Biolabs). The pellet was resuspended in 88 μL of NEBuffer 2.1 with 10 μL dNTP (1mM, #N0446S, New England Biolabs) and 2 μL T4 DNA Polymerase (#M0203S, New England Biolabs). The reaction was incubated at room temperature for 45 min with gentle mixing every 5 min. The sample was then washed 3X with 500 μL ice cold T4 DNA Ligase Reaction + 0.1% Triton X100 (#T8787, Sigma Aldrich). The pellet was then washed once with 500 μL T4 DNA Ligase Reaction Buffer. The pellet was resuspended in 18.5 μL T4 DNA Ligase Reaction Buffer with 5 μL of 10 μM annealed Broken end linker and 1.5 μL T4 DNA Ligase (#M0202S, New England Biolabs). The reaction was incubated 18-20 h at 16°C with constant mixing. The samples were washed 3X with T4 DNA Ligase Reaction + 0.1% Triton X100. The pellets were resuspended in 100 μL ChIP Buffer (SimpleChIP Kit #9003, Cell Signaling Technology), 2 μL RNAse A (10mg/mL, # EN0531, Thermo Fisher Scientific) and sonicated 30 sec on and 30 sec off for 45 min at 4°C. Samples were centrifuged at 835 x g for 10 min at 4°C, transferred to new tubes and frozen at −80°C. ChIP was performed using an amount of BEL-ChIP nuclear extract corresponding to 3.4 μg of input genomic DNA in 250 μL of ChIP Buffer + 1X PIC (SimpleChIP Kit #9003, Cell Signaling Technology). Antibody was added according to the vendor’s specifications and incubated overnight with end-over-end mixing at 4°C. Thirty μL magnetic protein G beads (SimpleChIP Kit #9003, Cell Signaling Technology) were added to the samples and incubated at 4°C with end-over-end rotation for 4 h. A magnetic rack was used to collect the beads, and the sample was washed 2X with 500 μL NEBuffer 2.1. The beads were resuspended in 100 μL of NEBuffer 2.1, 10 μL 1 mM dNTP, 2 μL T4 DNA Polymerase and incubated at room temperature for 25 min with constant shaking. The beads were placed in a magnetic rack to clear the supernatant and washed 2X with 500 μL ice cold T4 DNA Ligase Reaction Buffer. The beads were resuspended 50 μL T4 DNA Ligase Reaction Buffer with 5μL of 10 μM annealed Anchor linker and 1.5 μL T4 DNA Ligase. The sample was incubated overnight at room temperature with constant shaking. The beads were washed and DNA purified following the SimpleChIP Kit procedure. Purified DNA from the ChiP procedure (25 μL) was digested with 1 μL of I-SceI endonuclease (#R0694S, New England Biolabs) for 1 h and heat inactivated. PCR of 10 cycles with an annealing temperature 61°C was performed using Q5 High-Fidelity DNA Polymerase (#M0491S, New England Biolabs) and BEL and AL primers on 5μL of I-SceI digested DNA purified from the ChIP procedure. PCR reactions were run on a 2% agarose gel (NuSieve™ GTG™, Lonza) and the 200-400 bp region was excised and purified using a QIAquick Gel Extraction Kit (#28704, Qiagen). qPCR was performed using Power SYBR® Green PCR Master Mix (#4367659, Thermofisher Scientific), BEL and AL primers with annealing temperature of 61°C and 40 cycles on ABI 7900 system.

Broken End Linker

TACTACCTCGAGTAGGGATAACAGGGTAATTTTTTATTACCCTGTTTATCCCTACTCGAGGTAGTA

Anchor Linker CGTCGTCTCGAGTAGGGATAACAGGGTAATTTTTTATTACCCTGTTTATCCCTACTCGAGACGACG BEL Primer

TTATCCCTATCTCGAGGTAGTA

AL Primer

TTATCCCTACTCGAGACGACG

### Cell cycle staging and foci analysis

Cell cycle staging and foci quantification were performed on cells isolated from E10.5 *Dot1L^-/-^,* wild-type mice, HEK293T wild-type, *DOT1L^Stop^* and *DOT1L^Y312A^* cell lines using MANA (Machine Autonomous Nuclei Analyzer). Briefly, MANA was coded in Mathematica and utilized a convolutional (deep learning) neural network for nuclei classification. The neural network was trained using human curated nuclei images, and the data set was expanded to 180,000 nuclei images from both primary and tissue culture cells. Cell cycle staging was performed by using the Watson method for cell cycle peak calling (Watson, et al. 1987). Microscopy images were collected on a Nikon TE2000 Inverted microscope with a 40X objective, 0.6 NA.

### HR/NHEJ assays

Wild-type, *DOT1L^Stop^* and *DOT1L^Y312A^* HEK293T cell lines expressing an HR reporter (Pierce et al. 1999), alt-NHEJ reporter (Bennardo, et al. 2008), and a NHEJ reporter (Bennardo, et al. 2008), were transfected individually with Effectine (Qiagen) and selected using puromycin. Five days post-transfection, cells were transfected with the I-SceI plasmid using Effectine, and GFP-positive cells were counted 48 h later using an Invitrogen Attune flow cytometer. GFP values were normalized to the wild-type sample.

### Nucleosome stability assay

The nucleosome stability assay was performed essentially as described (Gursoy-Yuzugullu, et al. 2015). Briefly, wild-type, *DOT1L^Stop^* and *DOT1L^Y312A^* cell lines were exposed to 10 Gy IR and harvested at 30, 60, and 240 min post-IR. Cells were washed with ice-cold PBS and resuspended in 500 μL of Stability buffer (20 mM Hepes, pH 7.9, 0.5 mM DTT, 1.5 mM MgCl_2_ 2, 0.1% Triton) containing 1.0 M NaCl, Roche Protease Inhibitor cocktail) and agitated constantly for 40 min at 4 °C. Cells were collected by centrifugation at 100,000 × g (Beckmann Ultracentrifuge) for 20 min, the supernatant was removed, and the genomic DNA was resuspended in 100 μL ChIP Buffer (SimpleChIP Kit #9003, Cell Signaling Technology) with 2 μL RNAse A (10mg/mL, # EN0531, Thermo Fisher Scientific) and sonicated 30 sec on and 30 sec off for 45 min at 4°C. The DNA content was analyzed and all samples were adjusted to have 3.4 μg/mL DNA. A 1:10 dilution was made and 10 μL were analyzed by western blot.

### Comet assay and analysis

The alkaline comet assay for Fig. 1a was performed using the Trevigen Comet Assay kit (Trevigen, Gaithersburg, MD). Cells were isolated from yolk sacs of E10.5 *Dot1L^-/-^* and wild-type mice and grown in *ex vivo* differentiation media (see above). After 4 days cells were harvested and resuspended in cold PBS, and an aliquot of cells (1000/10 μl) was added to 100 μl of molten LMA agarose and spread onto a comet slide. The manufacturer’s protocol for alkaline comet assay was followed without modification. The slides were imaged using an upright Nikon Eclipse 80i fluorescence microscope at 20X, and analyzed using CometScore Pro (TriTek Corp). A neutralizing comet assay was used for monitoring the repair of induced DNA damage, Fig. 2a. Cells were isolated from E10.5 *Dot1L^-/-^* and wild-type mice and exposed to 2 Gy of ionizing radiation, and a neutralizing comet assay was performed as specified in the manufacturer’s protocol. Comets were analyzed using a custom comet analysis software development in Mathematica programming language.

### Statistical analysis

Sample size for each experiment is as follows: Figure 1A: WT mice, n > 100 cells; Dot1L^-/-^mice, n > 100 cells. Figure 1B: WT mice, n > 1000 cells; Dot1L^-/-^mice, n > 1000 cells. Figure 1C: WT mice, n = 3; Dot1L^-/-^mice, n = 3. Figure 1D: WT mice, n = 3; Dot1L^-/-^mice, n = 3. Figure 1E: WT mice, n > 1000 cells; Dot1L^-/-^mice, n > 1000 cells. Figure 2A: WT mice, n = 3; Dot1L^-/-^mice, n = 3. Figure 2B: WT mice, n = 3; Dot1L^-/-^mice, n = 3. Figures 3A and 3E: WT and Dot1L^-/-^mice, >250 cells. Figures 3B, 3C, 3D: WT HEK293T, *DOT1L^stop^* and *DOT1L^Y312A^* cells, n = 3. Figures 5A, 5B, 5C, 5D; WT HEK293T, *DOT1L^stop^* and *DOT1L^Y312A^* cells, n = 3, >250 cells each. Figures 5E and 5F: n > 1000 cells each for WT HEK293T, *DOT1L^stop^* and *DOT1L^Y312A^* cells. Figures 6B and 6C: n = 3 for WT HEK293T, *DOT1L^stop^* and *DOT1L^Y312A^* cells. Analysis was carried out in comparison to the control group. Two-sided t-tests were used for control vs. mutant in Figures 1C, 1D, 1E, 2A, 2B, 3B, 3C, 3D, and 6B. A Mann-Whitney test was used for control vs. mutant in Figures 1A, 3A, and 3E. A Z-Test was used for Figure 6C. Two-way ANOVA, Tukey’s multiple comparison tests were used for multiple comparison of groups in Figures 3A, 3B, 3C, and 3D.

## Code availability

Code used for MANA and comet analysis is available upon request.

