## Supplemental Data for "Dot1L-dependent H3K79 methylation facilitates histone variant H2A.Z exchange at DNA double strand breaks and is required for high fidelity, homology-directed DNA repair"

**Alvarez_Supplemental_Fig_1_legend**

**MANA (Machine Autonomous Nuclei Analyzer) pipeline overview.** High-resolution microscopy images of cells are segmented to isolate objects. The segmented objects are classified into two groups; nuclei and non-nuclei, by a deep learning neural network. Subsequent steps quantify foci numbers and cell cycle stage by calculating the intensity profiles of nuclei in 2D and 3D space.

**Alvarez_Supplemental_Fig_2_legend**

**Generation of *DOT1L*$^{STOP}$ and *DOT1L*$^{Y312A}$ mutant HEK 293 cells using CRISPR/Cas9.** (***A*** and ***B***) Sub-region of Human *DOT1L* locus with the CRISPR/Cas9 spacer sequence (blue) and PAM (orange). Nucleotides targeted for mutation are labeled in red. The region of interest in the single stranded oligonucleotide repair template is below. The added DdeI site is underlined in green. (**C**) PCR and DdeI resection digest screening strategy. (***D*** and ***E***) Subset of clones screened using PCR and DdeI digest to identify insertion. Positive clones are marked with asterisk. (***F*** and ***G***) Representative Sanger sequence of clones positive for mutation. (***H***) Example western blot of a positive clone.

**Alvarez_Supplemental_Fig_3_legend**

**H3K79 methylation deficient cells have normal G1 53BP1/BRCA1 foci ratio.** Mean of the ratio of 53BP1/BRCA1 foci in cells exposed to 10 Gy of IR in the G1 phase of cell cycle. Data are mean ± sem, n=3.

**Alvarez_Supplemental_Fig_4_legend**

**H3K79 methylation deficient cells have normal H4 lysine acetylation levels at double strand break sites.**

(***A***) Diagram of BEL-ChIP assay. (***B***) Histone stability assay following exposure to 10 Gy of IR in HEK293T cell lines. Western H4 acetylation. (***C***) BEL-ChIP of histone H4 lysine acetylation at sites of double strand breaks. Data are mean ± sem, n=3

**Alvarez_Supplemental_Fig_1**

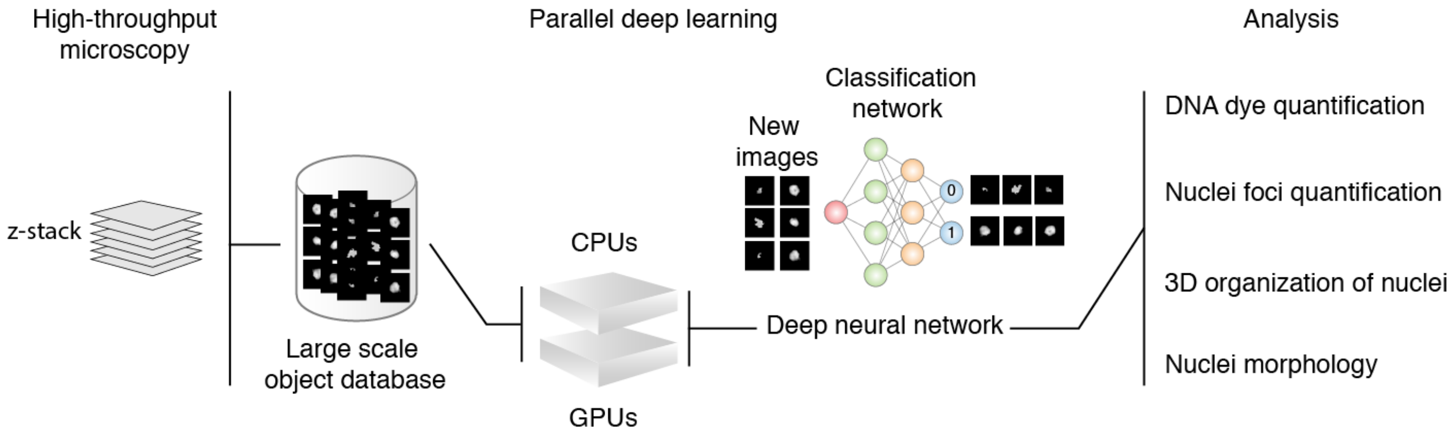

**Alvarez_Supplemental_Fig_2**

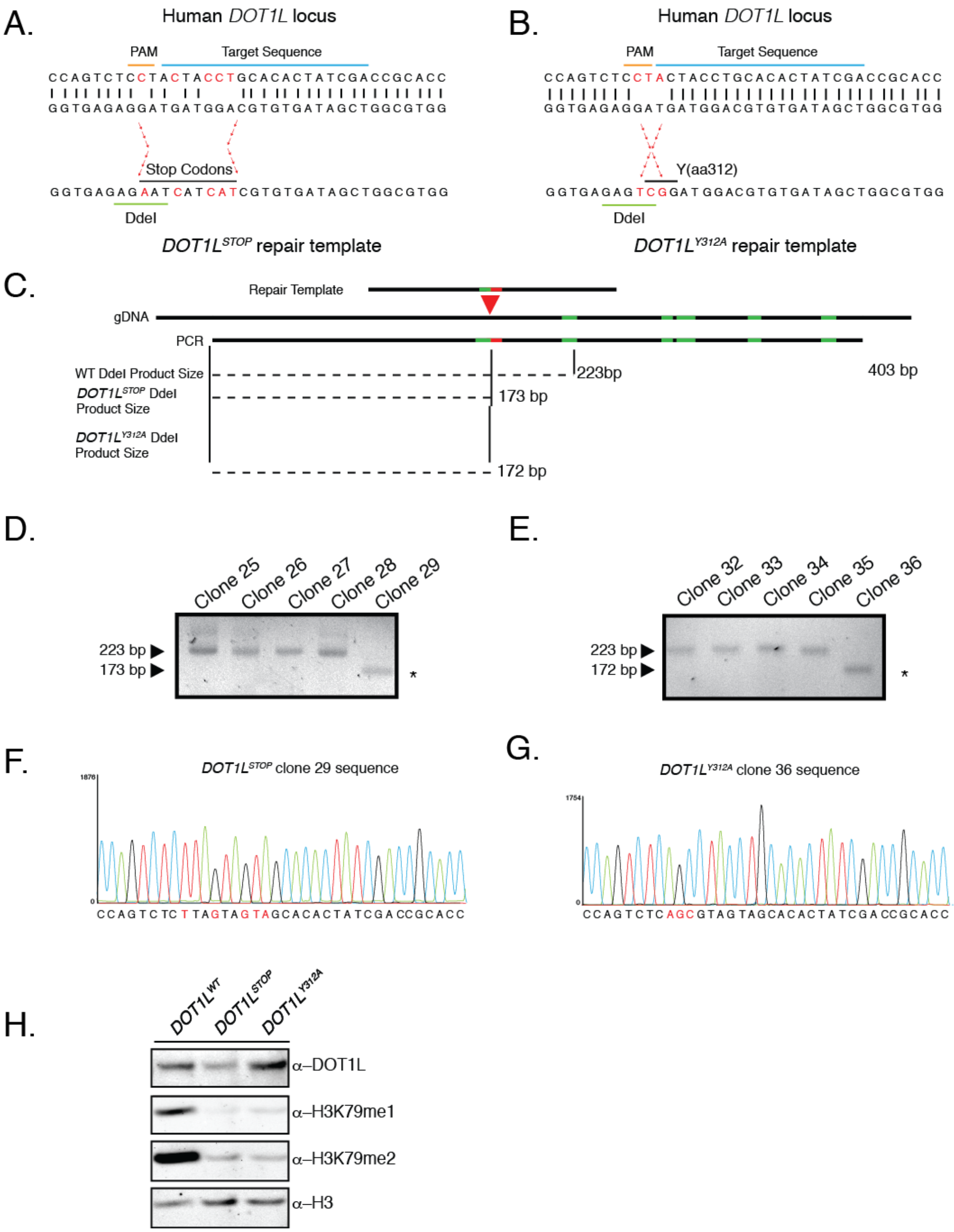

**Alvarez_Supplemental_Fig_3**

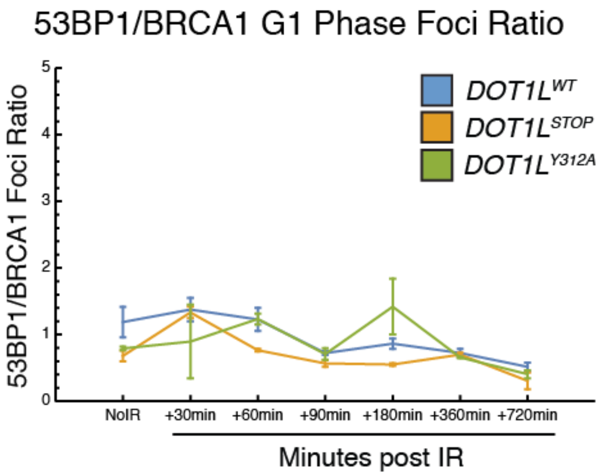

**Alvarez_Supplemental_Fig_4**

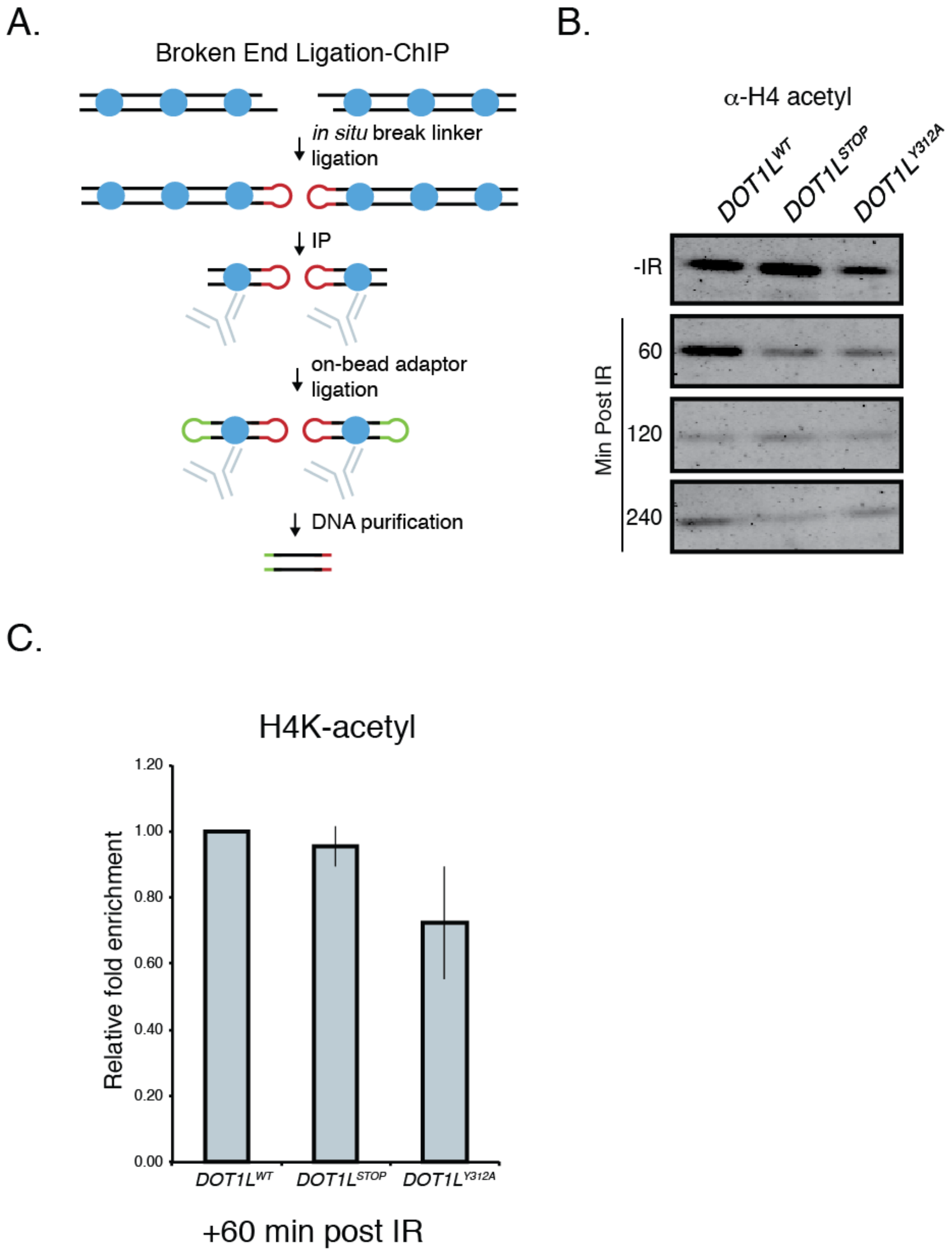
